# Novel agonist of GDNF family ligand receptor RET for the treatment of experimental neuropathy

**DOI:** 10.1101/061820

**Authors:** Maxim M. Bespalov, Yulia A. Sidorova, Ilida Suleymanova, James Thompson, Oleg Kambur, Viljami Jokinen, Tuomas Lilius, Gunnar Karelson, Laura Puusepp, Pekka Rauhala, Eija Kalso, Mati Karelson, Mart Saarma

## Abstract

Neuropathic pain is a chronic pain condition caused by lesion or disease affecting the somatosensory system. The glial cell line-derived neurotrophic factor (GDNF) family ligands (GFLs) alleviate symptoms of NP and stimulate regeneration of sensory neurons *in vivo*. Here we report the development of the compound BT18 that selectively activates GFL receptors, alleviates pain and restores damaged dorsal root ganglion (DRG) neurons in rat models of NP.

**Significance statement:** Neuropathic pain (NP) is a chronic syndrome caused by different diseases and lesions affecting nervous system. Earlier studies demonstrated that neurotrophic factors - the glial cell line-derived neurotrophic factor (GDNF) and artemin - could reverse the damage done by lesions in animal models of NP. We demonstrate for the first time that a small molecule can activate receptor of GDNF and artemin, it alleviates pain symptoms *in vivo* in two animal models of NP and restores to normal the molecular markers expressed in sensory neurons. This compound, termed BT18, can pave way for creating novel disease modifying therapies for NP.

## Introduction

Neuropathic pain (NP) affects millions of people yet existing therapies manage it poorly, do not remedy the underlying cause and have significant side effects. NP often accompanies physical trauma, chemotherapy treatment and diabetes (1–3).

GFLs are neurotrophic factors that support survival, regeneration and functioning of several neuronal populations: dopaminergic, enteric, sympathetic, parasympathetic, motor, cholinergic neurons and of sensory neurons. GFLs bind to the GDNF family receptor α(GFRα), which provides ligand-binding selectivity, and signal via a receptor tyrosine kinase (RTK) RET. GDNF preferentially binds to GFRα1 and ARTN to GFRα3 (Supplementary Fig. 1a). Ligand binding to GFRα/RET receptor complex stimulates RET phosphorylation and activation of intracellular signaling cascades including MAPK, PI3K/Akt, PLCγ, Src that are important for neuronal functionality (reviewed in (4); Supplementary Fig. 1b).

GDNF and artemin (ARTN) both can reverse neurological damage produced by lesion in animal models of NP (5–8). However, these proteins have poor pharmacological properties to be administered systemically (4, 9).

Using cell-based high-throughput screening we discovered novel family of compounds that could activate RET and induce its downstream signaling. We optimized the hit compounds to improve their potency an bioavailability and demonstrated that the compound termed BT18 could activate RET but no other tested RTKs *in vitro*, alleviated symptoms of NP *in vivo* and restored damaged DRG neurons in rat models of NP.

## Results

To identify small molecular weight GFL mimetics, we first made stable cell lines expressing GFL receptors and luciferase reporter that is induced by MAPK signaling (10) (Supplementary Fig. 1b). We screened 18000 diverse drug-like compounds using reporter cell line expressing GFRα1/RET, identifying 43 hits that increased luciferase activity at least three fold. We reconfirmed biological activity of five and further validated after re-synthesis two of them, both having a similar chemical structure, using the same luciferase-reporter assay. The hit-compound named BT13 (Supplementary Table 1; Fig. 1a) potently induced luciferase activity in reporter cell line, induced RET phosphorylation and RET-dependent intracellular signaling in GFRα1/RET-expressing cells (Supplementary Fig. 2) and it was chosen for future development. We found that BT13 also efficiently activated luciferase reporter in ARTN-responding GFRα3/RET-expressing cells (Fig. 1b) and less so in RET-only-expressing cells (Fig. 1a).

**Figure 1.**
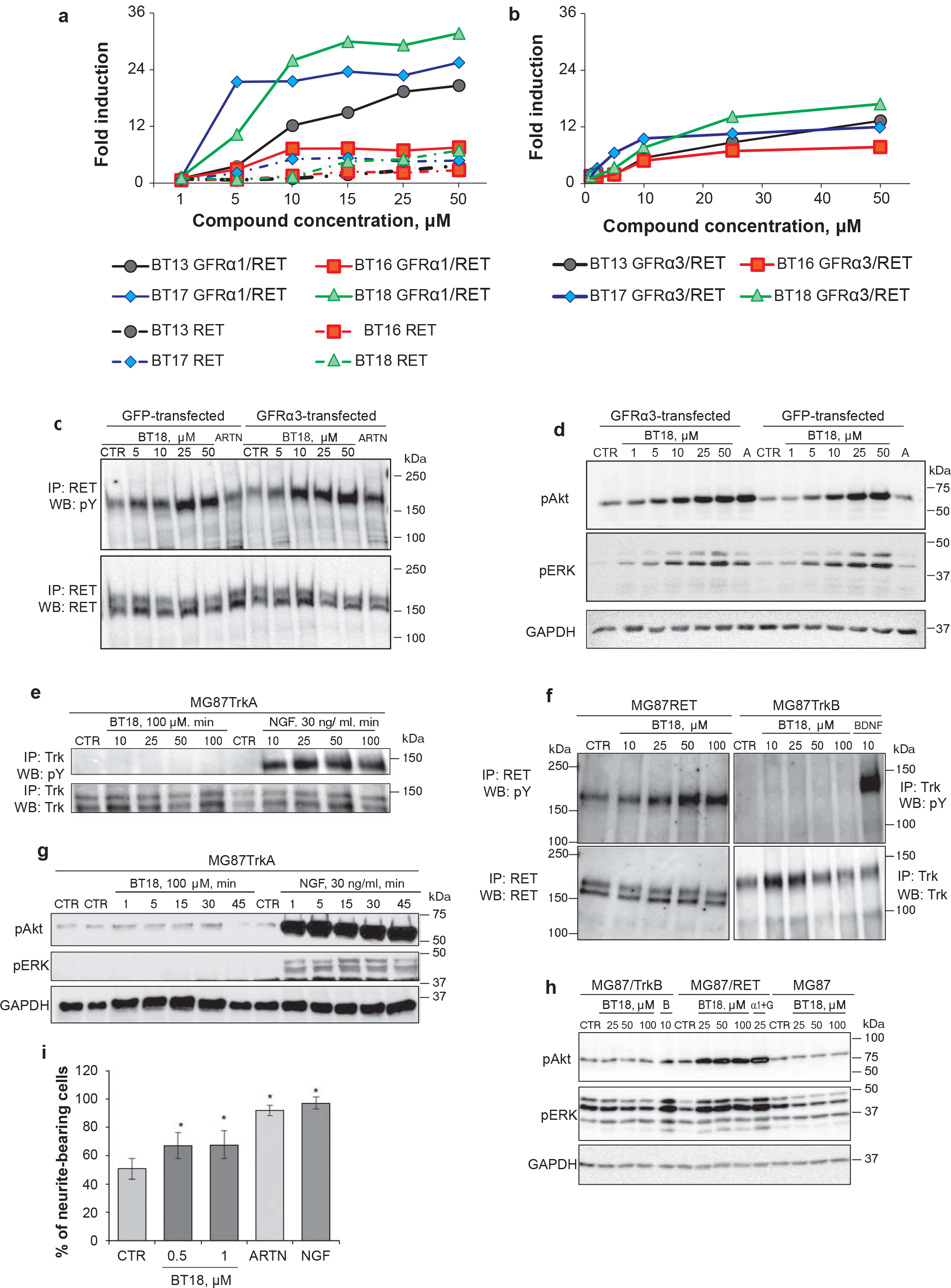
Design and development of small molecular weight RET agonists and their selectivity towards RET. (**a, b**) Dose-dependent activation of luciferase activity by BT13 and related compounds (BT16, BT17 and BT18) in RET-, GFRα1/RET-(**a**) and GFRα3/RET-expressing cells (**b**). (**c, d**) Dose-dependent activation of RET phosphorylation (**c**) and RET-dependent intracellular signaling (**d**) by BT18 in cells lacking GFRα3 (GFP-transfected) and expressing GFRα3 (GFRα3-transfected). (**e, f**) BT18 neither activates TrkA (**e**) nor TrkB (**f**) but does dose-dependently activate RET (**f**) in murine fibroblasts expressing corresponding receptors (MG87/TrkA, MG87/TrkB and MG87/RET). (**g, h**) Phosphorylation of Akt and ERK 42/44 in MG87, MG87RET, MG87TrkB cells in response to BT18, BDNF (B) and soluble GFRα1 with GDNF (α1+G). (**i**) Neurite outgrowth from mice neonatal DRG neurons. pY – phosphotyrosine, pAKT – phosphorylated form of AKT, pMAPK – phosphorylated forms of ERK 42/44, GAPDH – glyceraldehyde 3-phosphate dehydrogenase, loading control, IP – immunoprecipitation. CTR – control, A – ARTN, B – BDNF, G – GDNF. *p < 0.05 (ANOVA).

To improve the pharmacological properties of BT13 (effective concentration eliciting 50% of activation (EC_50_) in luciferase reporter assay in GFRα1/RET cells is 13μM) we synthesized compounds BT16, BT17 and BT18 with predicted improved activity towards GFRα/RET (Supplementary Table 1). We found that BT17 and BT18 indeed activated luciferase more potently than the original compound in both GFRα1/RET and GFRα3/RET lines (EC_50_ = 3 and 7 μM, in GFRα1/RET respectively) with almost no activity in RET-only expressing cells (Fig. 1a,b). All four compounds increased the level of RET phosphorylation in MG87RET fibroblasts expressing GFRα3 but also in cells lacking GFRα (Supplementary Fig. 3a). BT18 was the most active in RET phosphorylation assay in both cell lines and had insignificant acute toxicity to immortalized (Supplementary Fig. 3b) and primary murine fibroblasts (Supplementary Fig. 3c). We confirmed that BT18 dose-dependently increased phosphorylation of RET (Fig. 1c) and its downstream targets, Akt and ERK (Fig. 1d), in the presence and absence of overexpressed GFRα3 in MG87RET fibroblasts. We selected BT18 for further studies.

To elucidate the mechanism of BT18-induced RET activation we assessed direct binding of BT18 to GFL receptors. BT18 at 50 μM reduced binding of radioiodinated GDNF (^125^I-GDNF), which binds both GFRα1/RET and GFRα3/RET, in cells expressing GFRαl/RET ((Supplementary Fig. 4a), suggesting binding of GFLand mimetic to the same site. Since, GFRα appears to be dispensable for BT18 activity we attempted computational docking of the compound to RET at the site GDNF (and likely ARTN) interacts with RET (11). We found that BT18 can be docked to cysteine-rich domain (CRD) of RET (Supplementary Fig. 4b). In addition, RET downstream signaling can be induced by BT18 with fast kinetics, mimicking that of GDNF (Supplementary Fig. 4c). Together with ^125^I-GDNF displacement results and molecular docking this suggests a direct interaction of BT18 with RET, leading to productive RET signaling.

In addition to RET nociceptive sensory neurons express other RTKs such as neurotrophin receptors TrkA and TrkB (12). Since activation of TrkA and TrkB by their natural ligands, nerve growth factor (NGF) and brain-derived neurotrophic factor (BDNF), was shown to increase pain (13, 14), we tested the ability of BT18 to phosphorylate TrkA and TrkB and their downstream targets Akt and ERK (Fig. 1e-h) using MG87TrkA and MG87TrkB fibroblasts. We also assessed levels of pAkt and pERK upon BT18 administration in parental MG87 cells (Fig. 1h) lacking receptors for neurotrophic factor but expressing other RTKs, for example, fibroblast growth factor (FGF) receptors. BT18 neither activates TrkA (Fig. 1e), TrkB (Fig. 1f) nor ERK in the absence of RET (Fig. 1g,h). In addition, no activation of Akt was observed in MG87TrkB and MG87 parental cells in response to BT18 (Fig. 1h). However, in some experiments, BT18 elicited marginal and transient increase of pAkt level in MG87TrkA cells (Supplementary Fig. 5A). Taken together, our data indicate that BT18 is a selective RET agonist.

We then tested the ability of BT18 to mimic GFL activity in neurons and to stimulate neurite outgrowth from P1-P3 mouse DRG neurons. BT18 (0.5-1 μM) increased the number of neurite-bearing DRG neurons by approximately 30% *in vitro* (Fig. 1i, Supplementary Fig. 6).

We assessed **<u>a</u>**bsorption, **<u>d</u>**istribution, **<u>m</u>**etabolism and **<u>e</u>**xcretion (ADME) properties of BT18 (Supplementary Table 2) and considered the therapeutic window and the safety profile of BT18 satisfactory for testing in animal models of NP (Fig. 2a-e). Subcutaneous BT18 attenuated hyperalgesia/allodynia and increased thresholds of mechanical nociception of the ipsilateral paw in both the chronic constriction injury (CCI; Bennett, Fig. 2a,c,d) (15) and the spinal nerve ligation (SNL; Chung, Fig. 2a,c,e) (16) models of NP. The effect was statistically significant at doses above 5 mg/kg (Fig. 2e,d). Importantly, BT18 did not affect mechanical (Fig. 2f) sensitivity and motor performance (Fig. 2g) in non-neuropathic rats, indicating that its analgesic effect is specific for NP and not caused by sedation or impaired coordination. BT18 did not influence weight gain (Supplementary Fig. 7a), and no other signs of toxicity were observed during its administration.

**Figure 2.**
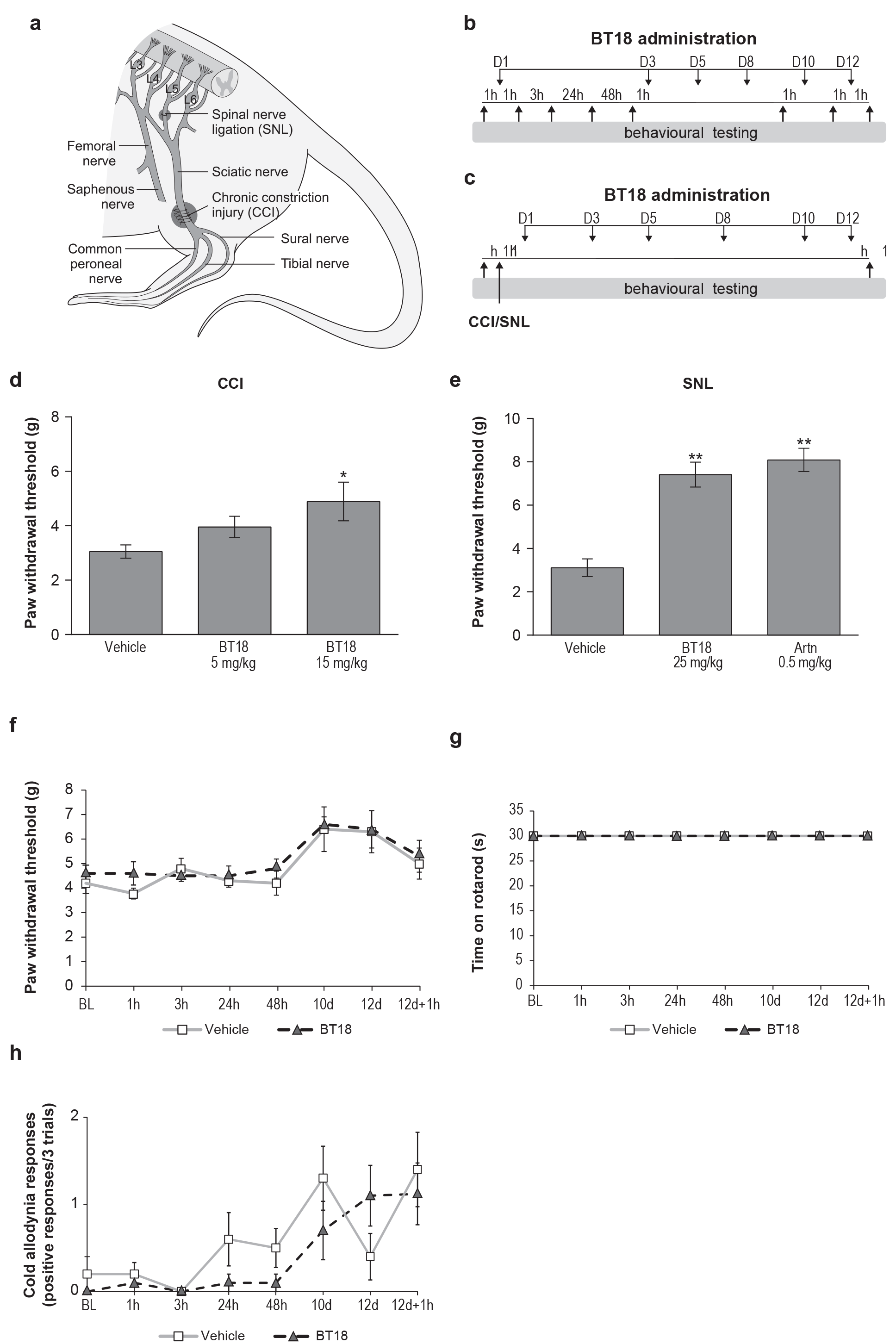
BT18 reduces mechanical hypersensitivity in animal models of neuropathic pain but does not affect sensitivity to mechanical or cold stimuli or motor coordination in non-neuropathic rats. (**a**) The scheme of neuropathic pain models used in this study: chronic constriction injury (CCI) and spinal nerve ligation (SNL). (**b, c**) The schedule of BT18 administration to rats without (**b**) and with (**c**) experimental neuropathy. (**d, e**) Thresholds of mechanical nociception (von Frey test) in animals with CCI (**d**) and SNL (**e**). (**f**) Responses of non-neuropathic rats treated with BT18 to mechanical (paw pressure test). (**g**) Motor performance (rotarod test) of rats treated with vehicle and BT18. (**h**) Responses of non-neuropathic rats treated with BT18 to cold stimuli (acetone test). In all experiments, groups were balanced on the basis of pre-treatment or pre-operative thr sholds (baseline, BL). The postoperative paw withdrawal thresholds of contralateral paws remained the same as BL and did not differ between groups in both CCI (BL, 44.8±3.0 g; postoperative, 42.2±3.2 g) and SNL (BL, 24.4±0.7 g; postoperative, 25.7±0.3 g) models of experimental neuropathy. D-days, *p < 0.05, **p < 0.0001 vs. the vehicle-treated group (ANOVA).

Since ARTN induced cold allodynia and thermal hyperalgesia in non-neuropathic rodents (17, 18), we assessed the effect of BT18 on thermal sensitivity in healthy rats, using acetone, hot-plate and tail-flick tests. BT18 neither provoked cold allodynia (Fig. 2h) nor induced tail-flick thermal hyperalgesia (Supplementary Fig. 7b), which might be explained by the selectivity of BT18 towards RET as the effect of ARTN protein on cold sensitivity is not RET-mediated (18). In the hot-plate test, however, BT18 produced some hyperalgesic effect (Supplementary Fig. 7c), as in earlier studies on ARTN (17, 18).

Similarly to ARTN (6), BT18 restored expression of different neuronal markers in DRGs (IB4, CGRP, NpY) and sciatic nerve (Nav1.8) of rats with SNL. In addition to what was shown for ARTN, we found that BT18 increased phosphorylation of ribosomal protein S6 (that is downstream target of PI3K/Akt pathway) and ERK (Fig. 3a-c) thus activating similarly to ARTN signaling cascades that support DRG neuron survival and axonal regeneration *in* vivo (4).

**Figure 3.**
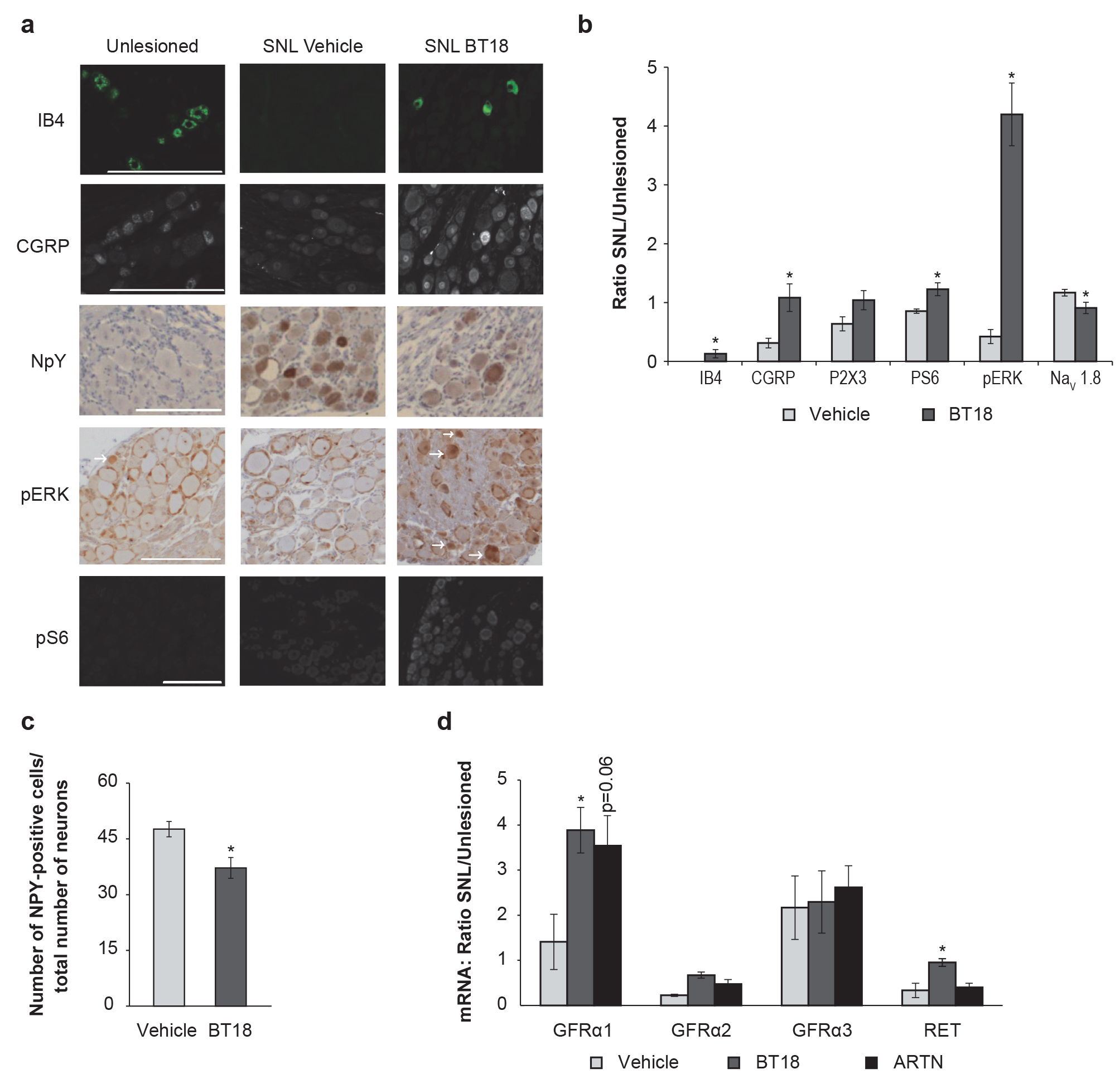
BT18 restores SNL-induced changes in expression of nociception-related neuronal markers, stimulates expression of GFLreceptors and activates GLF-like intracellular signaling. (**a**) Representative images of immunostained sections of DRGs. (**b, c**) Quantitative analysis of immunopositive cells on in DRGs. (**d**) qPCR analysis of GFRα1, GFRα2, GFRα3 and RET expression. We normalized the expression of specific marker on ligated side to unlesioned side to reduce variability whenever possible (**b, d**). Scale bar – 200 μm. SNL – spinal nerve ligation, * p < 0.05 by t-test (b, c), ANOVA with Tukey post hoc test (d).

We analyzed expression of RET, GFRα1, GFRα2 and GFRα3 mRNAs in DRGs of animals with SNL by qPCR since immunological methods using control tissues from respective knockout animals failed to produce reliable results. In line with earlier protein expression data (6), we observed an increase in GFRα3 mRNA level after SNL that was affected by neither ARTN nor BT18. However, in our study, SNL led to a marked decrease in RET expression that was restored by BT18 but not by ARTN. In addition, BT18 increased mRNA levels of GFRα1 but did not affect GFRα2 expression in DRGs of animals with experimental neuropathy (Fig. 3d).

## Discussion

Our results show that BT18 mimics the action of ARTN and GDNF *in vitro* and *in vivo* by activating RET. BT18 stimulates neurite outgrowth from cultured DRG neurons, alleviates NP manifestations in animal models and reverses surgery-induced changes in the expression of multiple neuronal markers in rat models of experimental neuropathy.

The microsomal stability of BT18 (t_1/2_ < 30 min) and its lifespan in serum (t_1/2_ = 41.7 min) are short (Supplementary Table 2). Therefore, *in vivo* BT18 is likely to activate RET transiently, which can prevent problems associated with chronic RET activation, such as possible tumor formation. In addition, BT18 has limited blood-brain permeability, reducing its potential for cerebral toxicity.

The mechanism of BT18 action warrants further study. Unlike natural ligands of RET (the GFLs), BT18 stimulates RET-dependent response even in the absence of co-receptor GFR**α**. In an active receptor complex the GDNF makes a direct contact with membrane-proximal domain of RET (11) and ARTN is likely to bind GFRα3/RET in the same manner. Our docking data indicate that BT18 can bind RET directly at the same site as do GDNF/ARTN (Supplementary Fig. 4). This may result in a conformational change seen in other RTK, leading to productive RET signaling (4). The hypothesis suggesting the direct interaction of BT18 and RET is supported by kinetics of RET activation by BT18 and by the displacing of radiolabeled GDNF from cells expressing GFRα1/RET by BT18 (Supplementary Fig. 4). Further biochemical experiments with labelled ARTN and purified components of the GFRα3/RET complex are required to elucidate the exact mechanism of BT18 action.

The previously described compound, XIB4035 (20), modulating GFRα/RET signaling and alleviating pain symptoms in small-fiber neuropathy animal model could not activate RET in the absence of GFLand is, therefore, could not be considered as RET agonist.

In summary, BT18 is the first described RET small molecule agonist that significantly attenuates pain behavior, restores molecular changes caused by experimental nerve injury in peripheral neurons *in vivo* and thus opens a way to develop novel disease-modifying drugs for neuropathic pain treatment.

## Materials and methods

**Animal.** Animal experiments were conducted in accordance to local and European regulation, guidelines of the International Association for the Study of Pain (19) and guided by the 3R principle. The provincial government of Southern Finland (Etelä-Suomen aluehallintovirasto, Hämeenlinna, Finland, ESAVI/5684/04.10.03/2011) approved the *in vivo* study concept regarding the behavioral analysis. The use of experimental animals for isolation of primary neurons was approved by the Committee for Animal Experiments of the University of Helsinki, and the chief veterinarian of the County Administrative Board permission (KEK11-025, VKL011-08). Primary DRG neurons were isolated from P1-P3 NMRI mice.

Acute pain sensitivity and locomotor activity were tested on male Wistar Han rats (Harlan, Netherlands), weighing 190-250 g, at the University of Helsinki. Experiments in animal models of neuropathic pain were ordered from Psychogenics Inc. (USA) and performed on male Sprague Dawley rats (100-125 g) from Harlan (Indianapolis, IN). The pharmacokinetic study was carried out by Cyathus Exquirere (Milan, Italy) on male Sprague Dawley rats (150-175 gr) supplied by Harlan, Netherlands. In all cases the animals were housed in groups (n = 3-5/cage) at ambient temperatures of 19-25°C. During the course of the study, 12/12 light/dark cycles were maintained, water and standard laboratory chow were provided *ad libitum*.

**Models of neuropathic pain.** Neuropathic pain in rats was initiated by chronic constriction injury of the left sciatic nerve (CCI, Bennett’s model of neuropathic pain (15)) or by ligation of the left L5 spinal nerve (SNL, Chung’s model of neuropathic pain (16)). Before operation, the rats were tested for sensitivity to mechanical stimuli using von Frey (VF) filaments. Animals with a paw withdrawal threshold (PWT) below 12 g were excluded from the study. Rats were subsequently balanced and assigned to treatment groups based on their pre-operative PWT values. Surgery was performed under isoflurane anesthesia. All rats received an analgesic (buprenorphine, 0.05 mg/kg, subcutaneously) immediately before and 6 h after surgery. Each rat was monitored until it was awake and moving freely around the recovery chamber. Animals were then single-housed for the duration of the study.

**Treatments.** Animals received 5, 15 and 25 mg/kg of BT18 dissolved in corn oil containing 5% DMSO or 0.5 mg/kg of ARTN protein dissolved in saline or vehicle, subcutaneously on the 1^st^, 3^rd^, 5^th^, 8^th^, 10^th^ and 12^th^ day of the experiment. Responsiveness of animals with neuropathic pain to the standard analgesic for neuropathic pain, gabapentin, was confirmed using a single dose of gabapentin (100 mg/kg, orally, 1 h prior to behavioral tests). The treatment regimen (except gabapentin) was started 1 h postoperatively on the day of surgery. On the 12^th^ day of the treatment, the treatments were administered to the rats 1 h prior to behavioral studies.

**Behavioral tests.** Sensitivity of the rats with experimental neuropathy to mechanical stimuli was determined using VF filaments on the 12^th^ day after surgery. VF filaments of increasing bending force were applied to the plantar surface of the hind paw. Pressure required to elicit withdrawal of the paw was determined.

In non-neuropathic rats, we assessed cold sensitivity, acute nociception and motor coordination using the acetone test (AC), tail-flick (TF), hot plate (HP), paw pressure (PP), and rotarod (RR) apparatuses (Ugo Basile, Comerio, Italy, device models 37360, 35100, 37215, and 47700, respectively). We measured baseline responses on the 1^st^ day of the experiment, 1 h before the administration of the test compounds. Consequently, animals were divided into two balanced groups according to weight and nociceptive baselines. The experiments were performed in a randomized and blinded fashion.

To measure sensitivity to thermal stimuli, we used TF and HP tests. For the TF test, the infrared light was directed in turn to three different points of the middle third of the tail of a constrained animal. For the HP test, the rat was put on a hot plate adjusted to 52°C (± 0.2°C). Latency time before the TF or licking or brisk shaking of the hind paw or jumping was assessed and cut-off times of 7 TF and 60 s HP were used to prevent tissue damage. Sensitivity to mechanical stimuli was measured with the PP test. The left hind paw of the constrained rat was placed under a pivot and the force applied to the paw was linearly increased. Vocalization or brisk shaking of the hind limb was used as a behavioral sign of nociception and the measurement was terminated. The force causing the behavioral response was recorded. The cut-off was set to 500 g. To assess motor coordination, the time during which the animal was able to stay on the rotating rod (20 rpm, fixed speed) was measured and a cut-off time of 30 s was used. Cold allodynia was assessed using the AC (18). The animals were placed in an elevated plexiglass chamber with a mesh floor and allowed to acclimatize for 5 min. A drop of acetone was applied to the plantar surface of the hind paw three times with 3-min intervals between the measurements, alternating paws between stimulations. A brisk withdrawal, shaking or licking of the hind limb were considered as responses of cold allodynia and their number was recorded.

**Phosphorylation assays.** Phosphorylation of receptor tyrosine kinase RET, TrkA, TrkB in cultured cells was assessed by antibodies against phosphorylated tyrosine residues (clone 4G10, Upstate) after immunoprecipitation of RTKs with specific antibodies targeted to RET or TrkA/TrkB, respectively. Phosphorylation of intracellular signaling proteins ERK and AKT was determined using specific antibodies against phosphorylated forms of the respective proteins.

**Cytotoxicity assays.** To assess the ability of our compounds to influence proliferation of immortalized (MG87 murine fibroblasts) and primary non-neuronal cells (embryonic fibroblasts), we used AlamarBlue dye (Life Technologies) or CellTiterGlo (Promega).

^125^**I-GDNF displacement assay.** GDNF was enzymatically iodinated by lactoperoxidase as described (21). Cells were treated with 5-100 μM BT18 or 5 nM unlabelled GDNF and 50 pM of ^125^I-GDNF dissolved in blocking solution and incubated for 2 h on ice, washed and lysed with 1 M NaOH. Lysates were transferred to vials and counted on a Wallac Gamma Counter (Wallac/LKB).

**ADME-Tox profiling.** ADME profiling was performed by CRO Cyathus Exquirere (Milan, Italy). To assess BT18 pharmacokinetics, 10 mg/kg of BT18 were intravenously injected into nine male Sprague Dawley rats. Plasma and brain samples were collected 1, 3 and 6 h later.

**Immunohistochemistry.** Sections of sciatic nerves and DRGs from rats with spinal nerve ligation and treated with BT18 or vehicle were probed, after deparaffinization and acidic or basic antigene-retrieval using standard IHC protocols, with antibodies against NaV1.8 (1:1000, Abcam, UK), IB-4 (1:200, Griffonia), P2X3 (1:1000, Neuromics), CGRP (1:10000, Peninsula Laboratory), NPY (1:10000, Peninsula Laboratory), pERK1/2 (1:300, Cell Signaling), pS6 (1:300, Cell Signaling), pan-neuronal marker PGP9.5 (1:500 Abcam) and nuclear dye.

**qPCR.** Expression of GFL receptors (GFR**α**1-3, RET) in DRGs of animals with SNL receiving vehicle, BT18 or ARTN (0.5 mg/kg) were analyzed by qPCR on LightCycler^®^ 480 SYBR Green I Master Mix (Roche Diagnostics). Statistical analysis was performed using data from 4-5 individual animals per group.

**Statistical analysis.** All quantitative data were subjected to statistical analysis. Results are presented as M±m, where M is average and m is standard deviation or standard error of mean. Significance of the difference between treatment groups was determined by Student’s *t*-test (two groups or paired values) or ANOVA (multiple groups), followed by Fisher’s or Tukey post-hoc comparisons, if criteria of parametric analysis were met. To analyze non-parametric data (acetone test results), we used the Mann-Whitney U-test.

Details for all methods are available in *SI Methods*.

## Acknowledgements

Current work was financially supported by FP7-HEALTH-2013-INNOVATION-1 GA N602919, FP7-PEOPLE-2013-IAPP GA N612275, CIMO, Genecode Ltd, Chemedest Ltd., Lundbeck Foundation, Sigrid Jusélius Foundation, EU European Regional Development Fund through the Center of Excellence in Chemical Biology, Estonia and by targeted financing from the Estonian Ministry of Education and Research (SF0140031As09). We acknowledge Psychogenics Inc (USA) for the assessment of BT18 activity in rat models of neuropathic pain and CRO Cyathus Exquirere (Italy) for ADMET studies. We thank Dr. Tomi Rantamaki and Hanna Anttilla for advice on TrkB phosphorylation experiments. Dr Pia Runeberg-Roos is acknowledged for help with RET phosphorylation assay. We are grateful to Laura Salminen, Sascha Gromnitza, Elisa Piranen, Jenni Montonen for assistance with the experimental procedures. Drs Urmas Arumäe, Mikko Airavaara and Jaan-Olle Andressoo are acknowledged for the critical comments on the manuscript. We are grateful to Les Hearn and MSc. Katrina Albert for proofreading.

## Supplemental Information

**Supplementary Figure 1.**
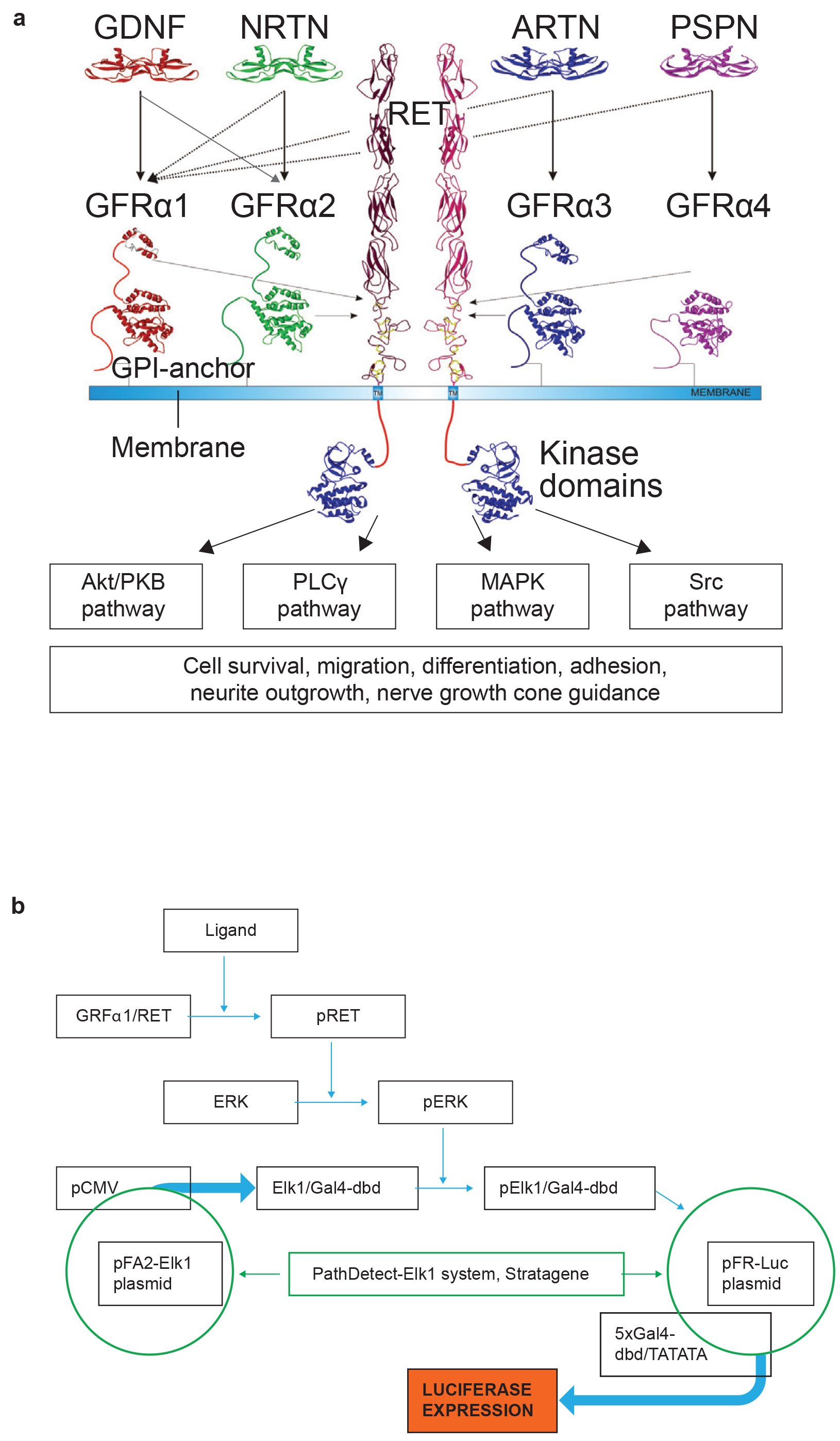
Scheme of GFL signaling and downstream events (a) and a schematic diagram of cell-based luciferase-reporter high-throughput assay for GFL mimetics (b). (a) Stabilization of GFRα/RET complex by GFLs (GDNF, NRTN, ARTN and PSPN) leads to kinase activation and downstream signaling pathways activation. (b) Diagram representing molecular mechanisms underlying an assay for detecting RET agonists. RET activation leads to MAPK signaling pathway activation. Activated forms of ERK 42/44 (pERK) phosphorylate transactivation domain of a fusion protein consisting of Elk1 transcription factor and yeast Gal4-DNA binding domain. Fusion protein is controlled by CMV promotor (pCMV) and constitutively expressed by PFA-Elk1 plasmid from PathDetect Elk1 system (Stratagene). Phosphorylation of the fusion proteins leads to its interaction with Gal4 sequence that controls expression of luciferase in pFR-Luc plasmid (Stratagene). NRTN – neurturin, PSPN – persephin, pRET, pERK, pElk1/Gal4-dbd – phosphorylated forms of corresponding proteins; dbd - DNA binding domain.

**Supplementary Figure 2.**
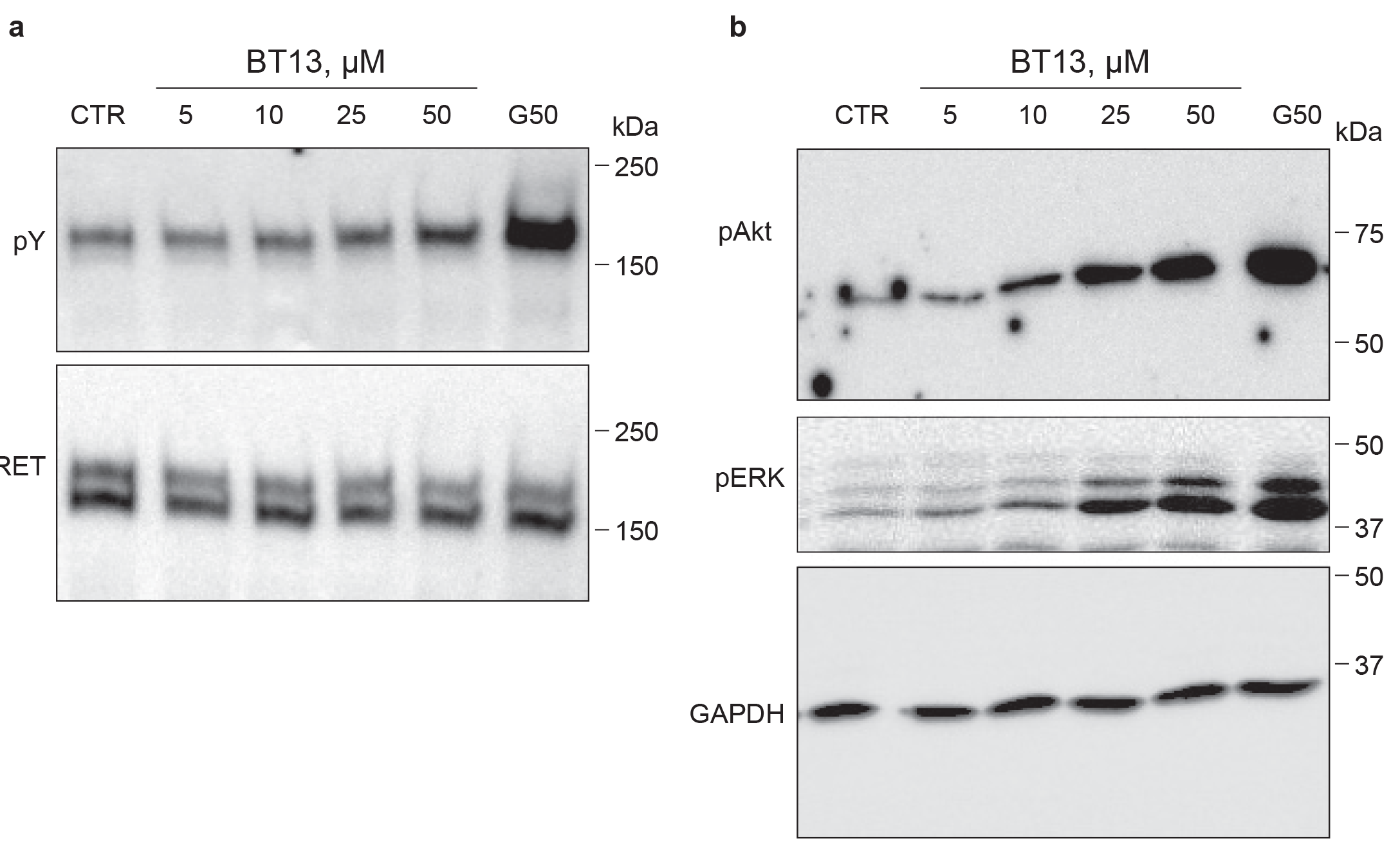
BT13 activates RET and RET-dependent signaling as determined by protein phosphorylation assays. Dose-dependence for BT13-induced phosphorylation of tyrosine residues (pY) in RET (a) and phosphorylation of RET downstream targets Akt and ERK (b). G50 – GDNF, 50 ng/ml.

**Supplementary Figure 3.**
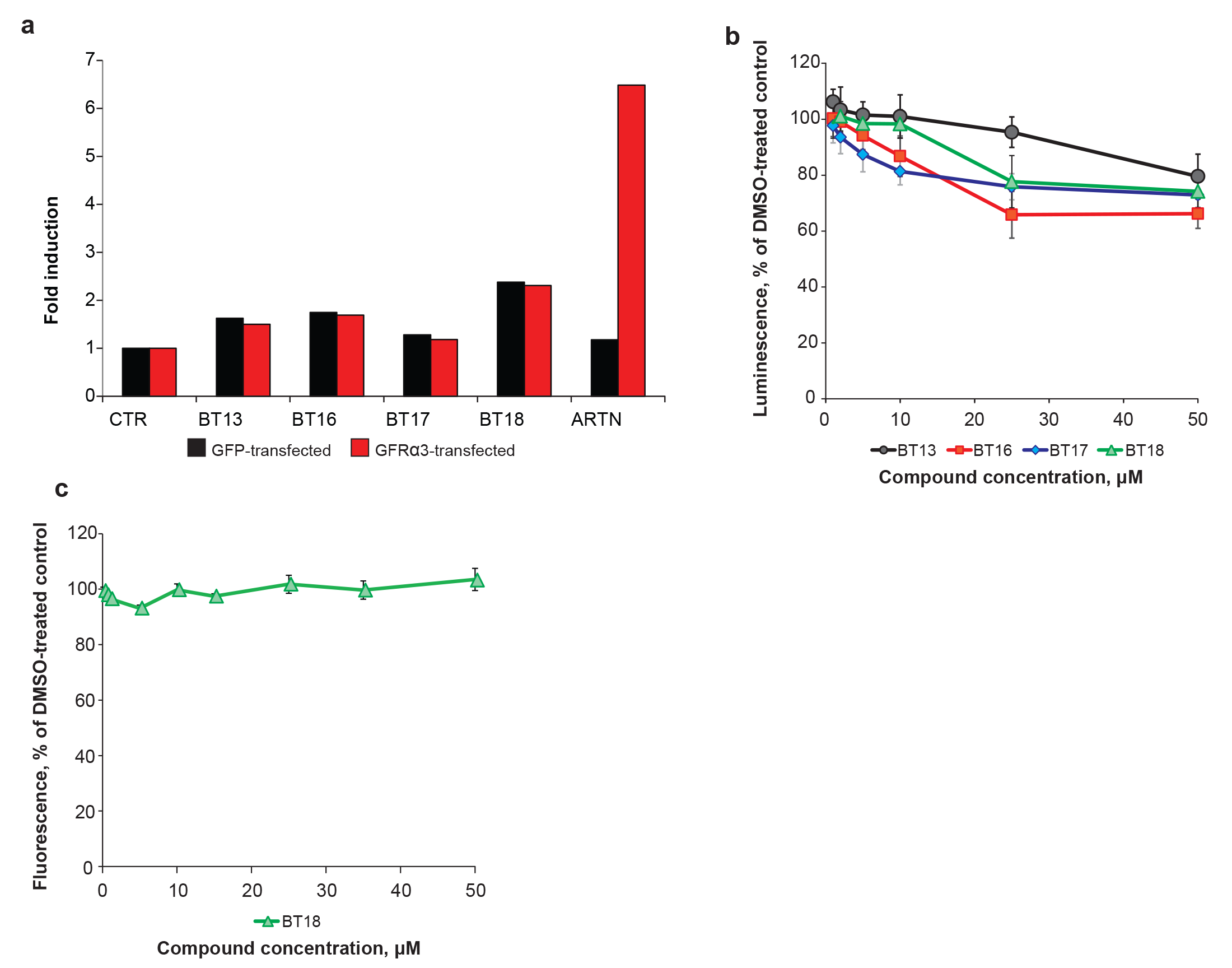
Effects of GFL mimetics on RET phosphorylation (a) and cell proliferation (b, c). (a) RET phosphorylation in response to GFL mimetics (50 μM) in GFP- and GFRα3-transfected MG87RET fibroblasts determined by RET-ELISA method (a); influence of GFL mimetics on proliferation of MG87 cells (b) and of BT18 on primary embryonic fibroblasts growth (c).

**Supplementary Figure 4.**
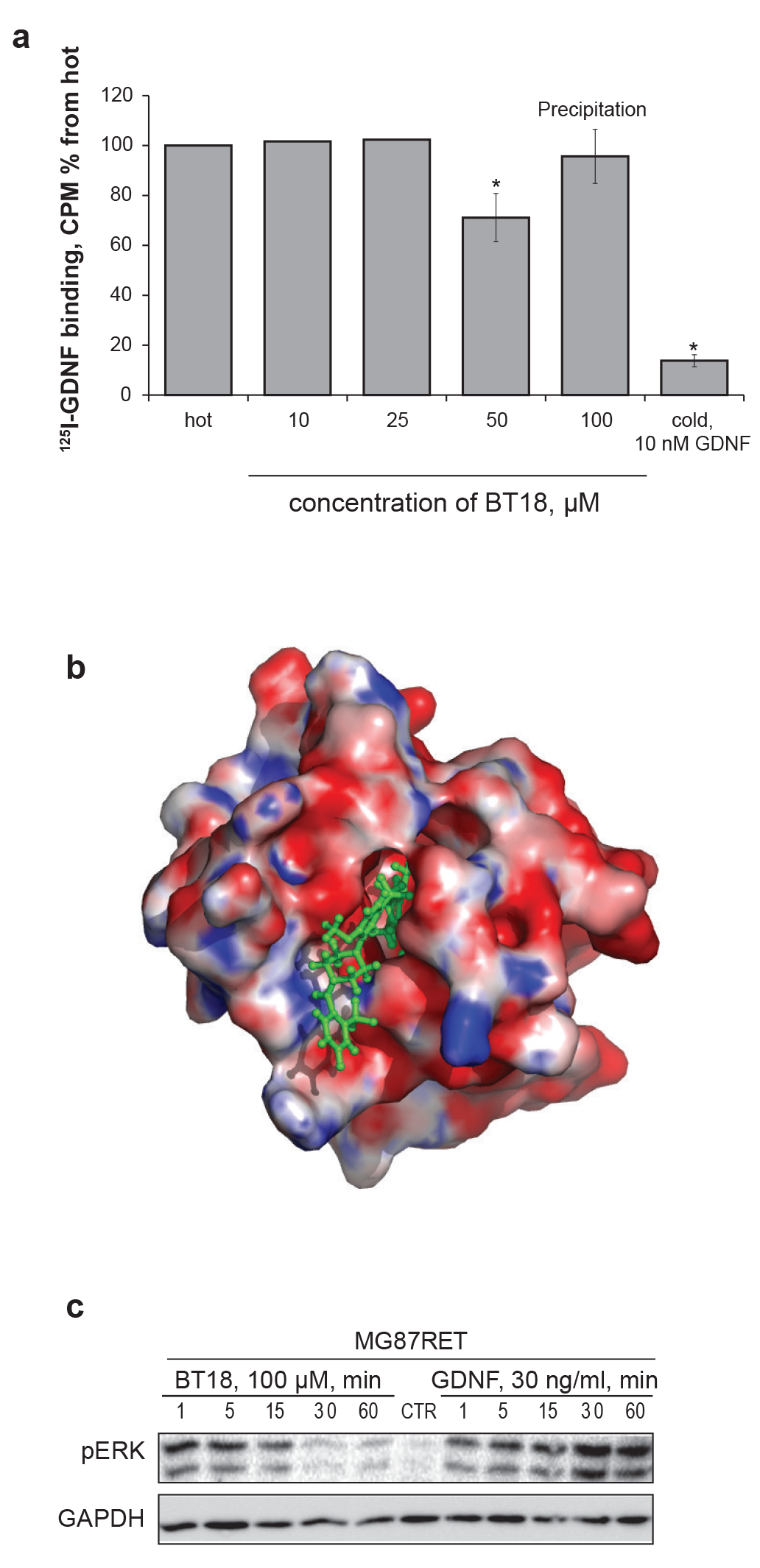
Displacement of 125I-GDNF by BT18 in cells expressing GFRμ1/RET (a). (b) Docking of BT18 to the homology model of RET cysteine rich domain. BT18 is represented as green ball-and-stick model. Protein is represented as electrostatic potential surface map. Red color represents negatively charged areas, blue color marks positively charged regions (b). (c) Time-dependent activation of intracellular ERK signaling by BT18 and GDNF in RET-expressing fibroblasts (MG87RET). CPM – counts per minute, Hot – iodinated GDNF, Cold – iodinated GDNF together with 500-fold molar excess of unlabeled GDNF. Precipitation – at 100 μM we observed BT18 precipitating in the binding buffer. GAPDH – glyceraldehyde-3-phosphate dehydrogenase. pERK – phosphorylated form of ERK. * p < 0.05, paired one-tailed t-test.

**Supplementary Figure 5.**
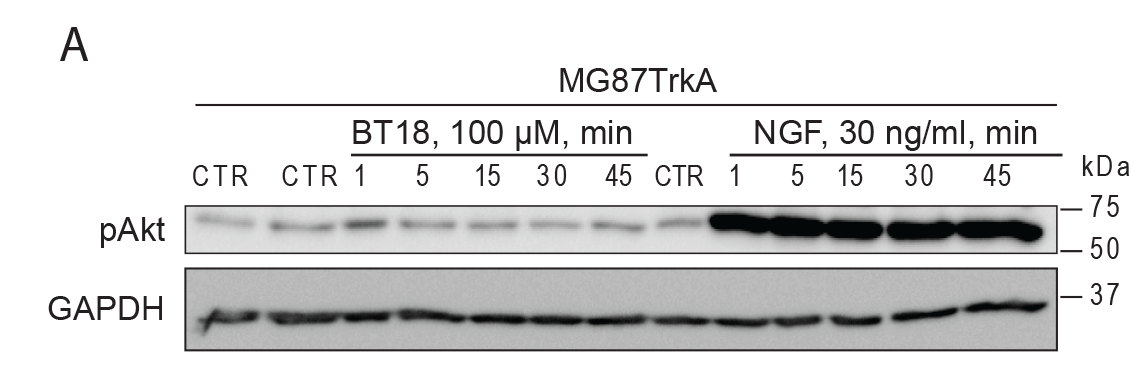
Time-dependent activation of intracellular signaling cascades in MG87TrkA (a). Only marginal and temporal activation of Akt phosphorylation in MG87TrkA cells observed in several experiments. pAKT – phosphorylated form of AKT, pERK – phosphorylated form of ERK, GAPDH – glyceraldehyde-3-phosphate dehydrogenase.

**Supplementary Figure 6.**
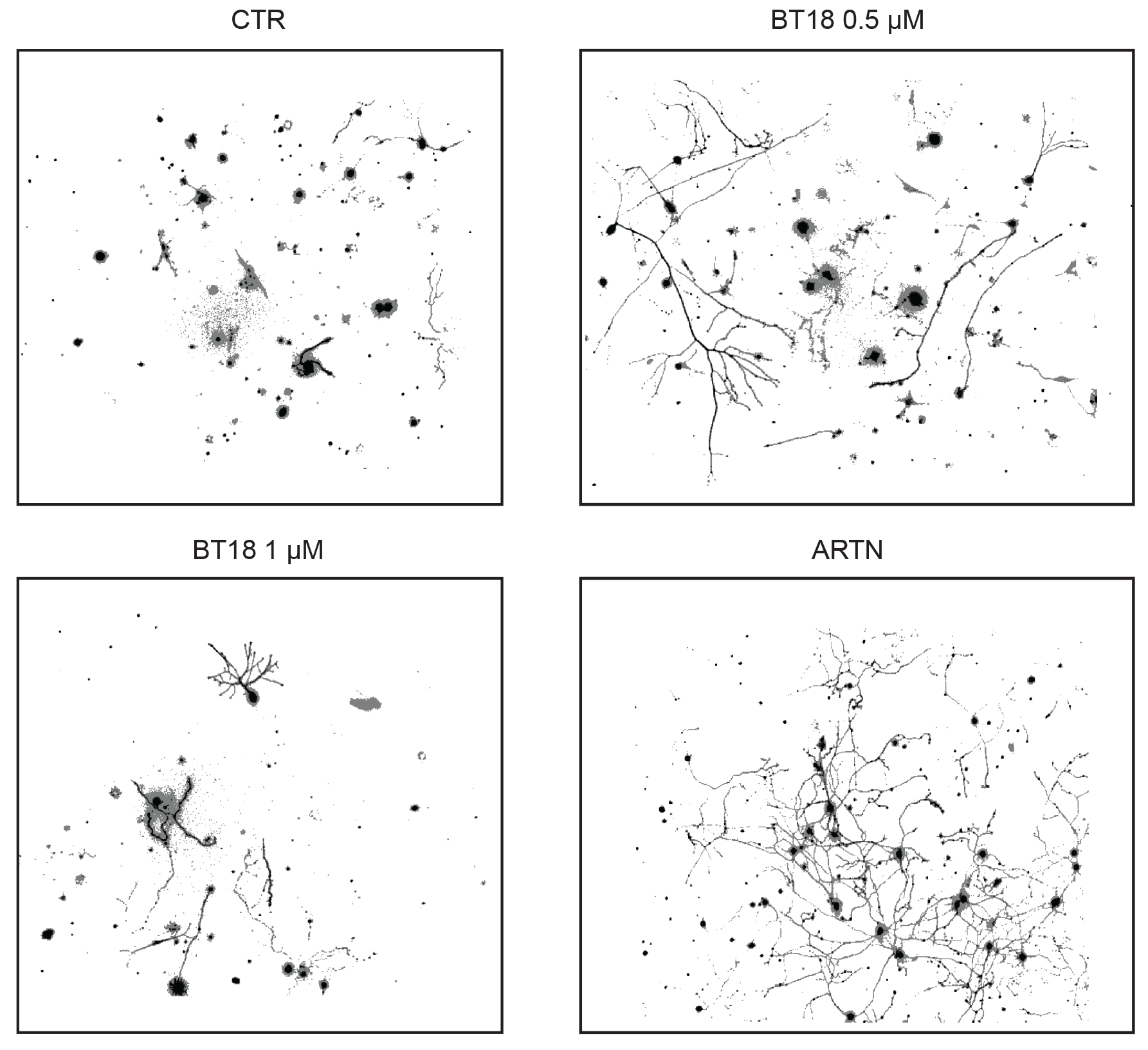
Neurite outgrowth from cultured P1 mouse DRG neurons treated with vehicle, BT18 or ARTN.

**Supplementary Figure 7.**
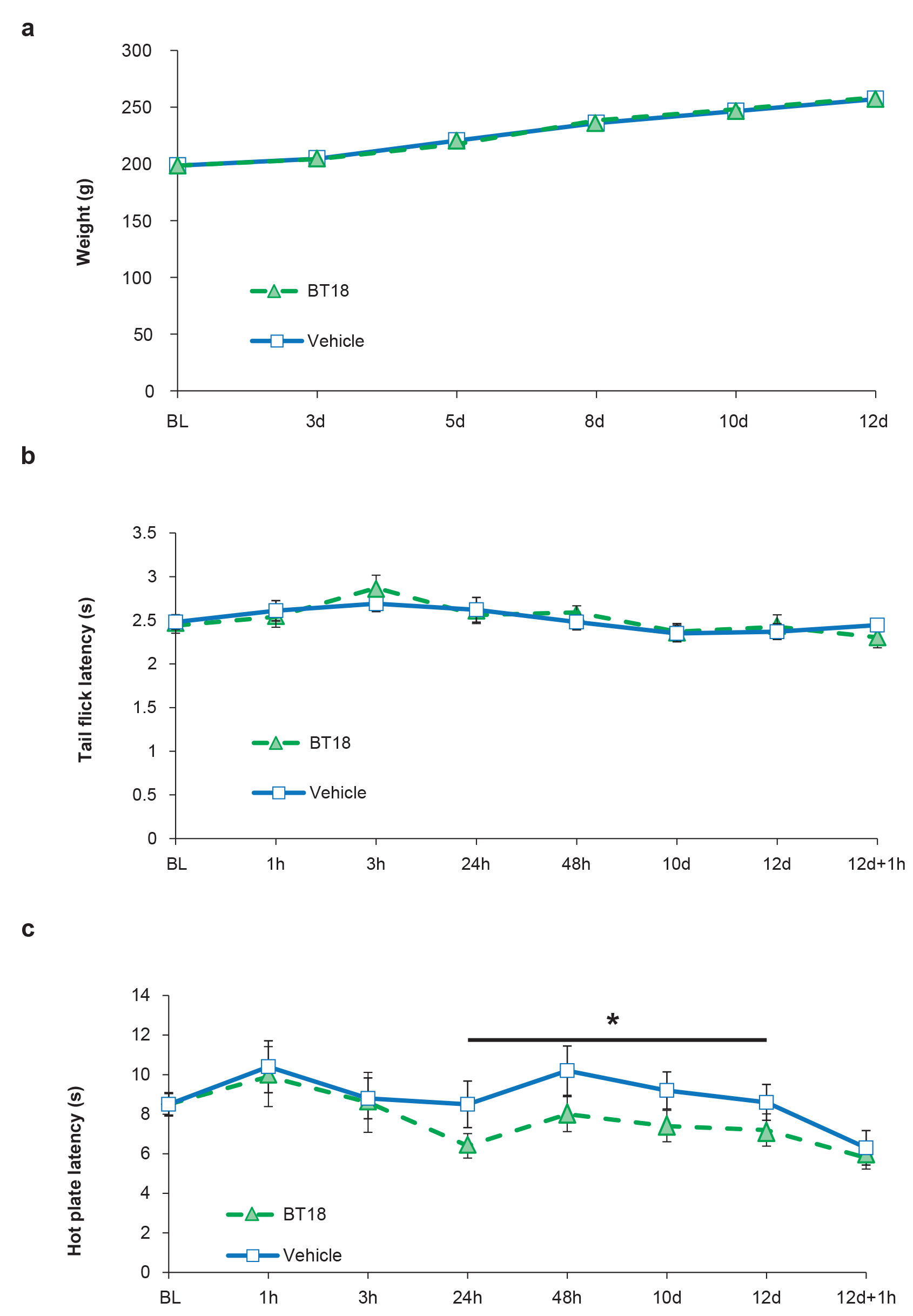
Effect of repeated administration of BT18 (25 mg/kg, s.c.) or vehicle on weight development (a) and thermal nociception assessed with tail flick (b) and hot plate (52 ± 0.2°C) (c) of male Wistar Han rats (n = 10/group). * p < 0.05 vs vehicle-treated, ANOVA.

**Supplementary table 1.**
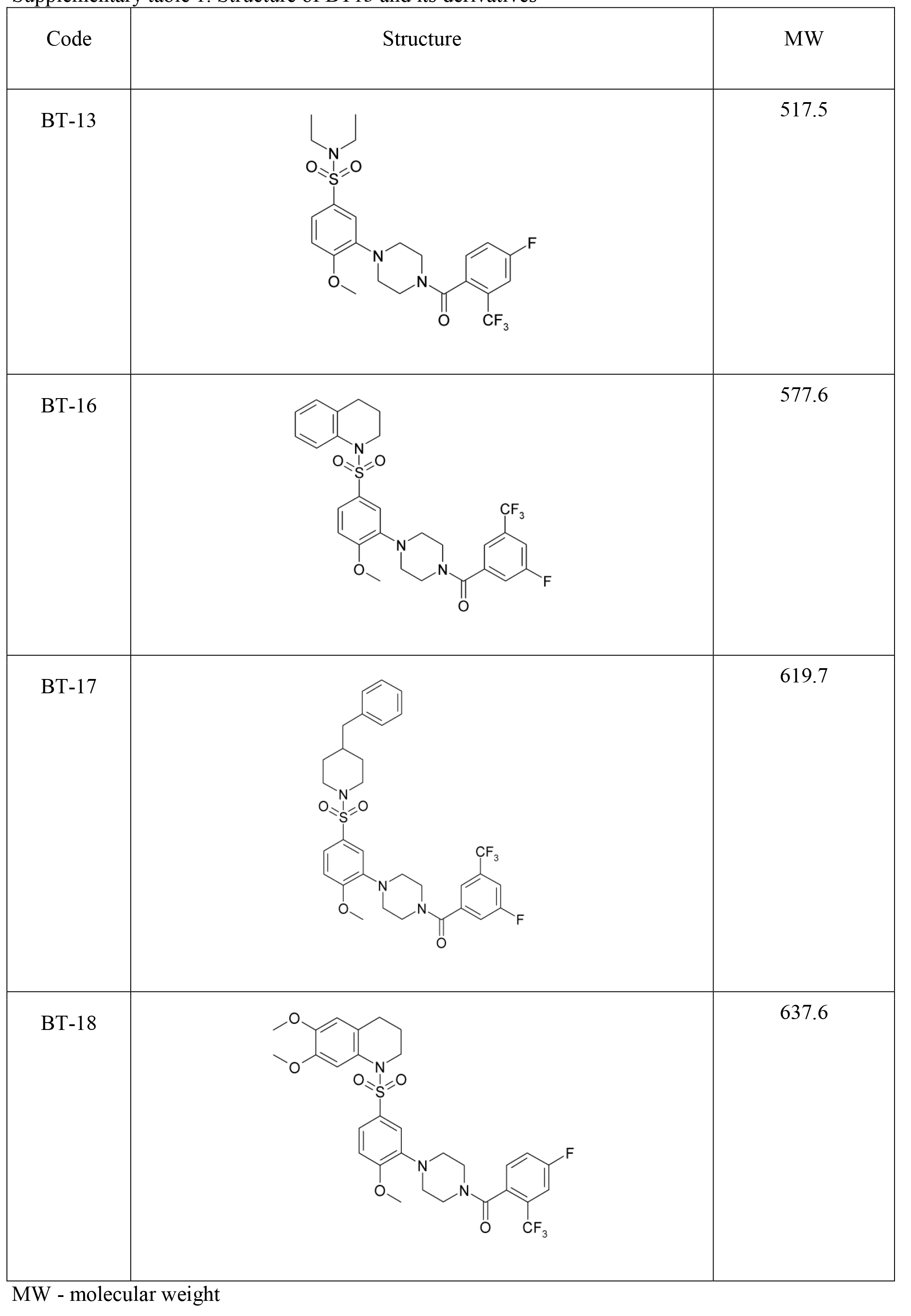
Structure of BT13 and its derivatives.

| Code | Structure | MW |
| --- | --- | --- |
| BT-13 |  | 517.5 |
| BT-16 |  | 577.6 |
| BT-17 |  | 619.7 |
| BT-18 |  | 637.6 |

MW - molecular weight

**Supplementary table 2.**
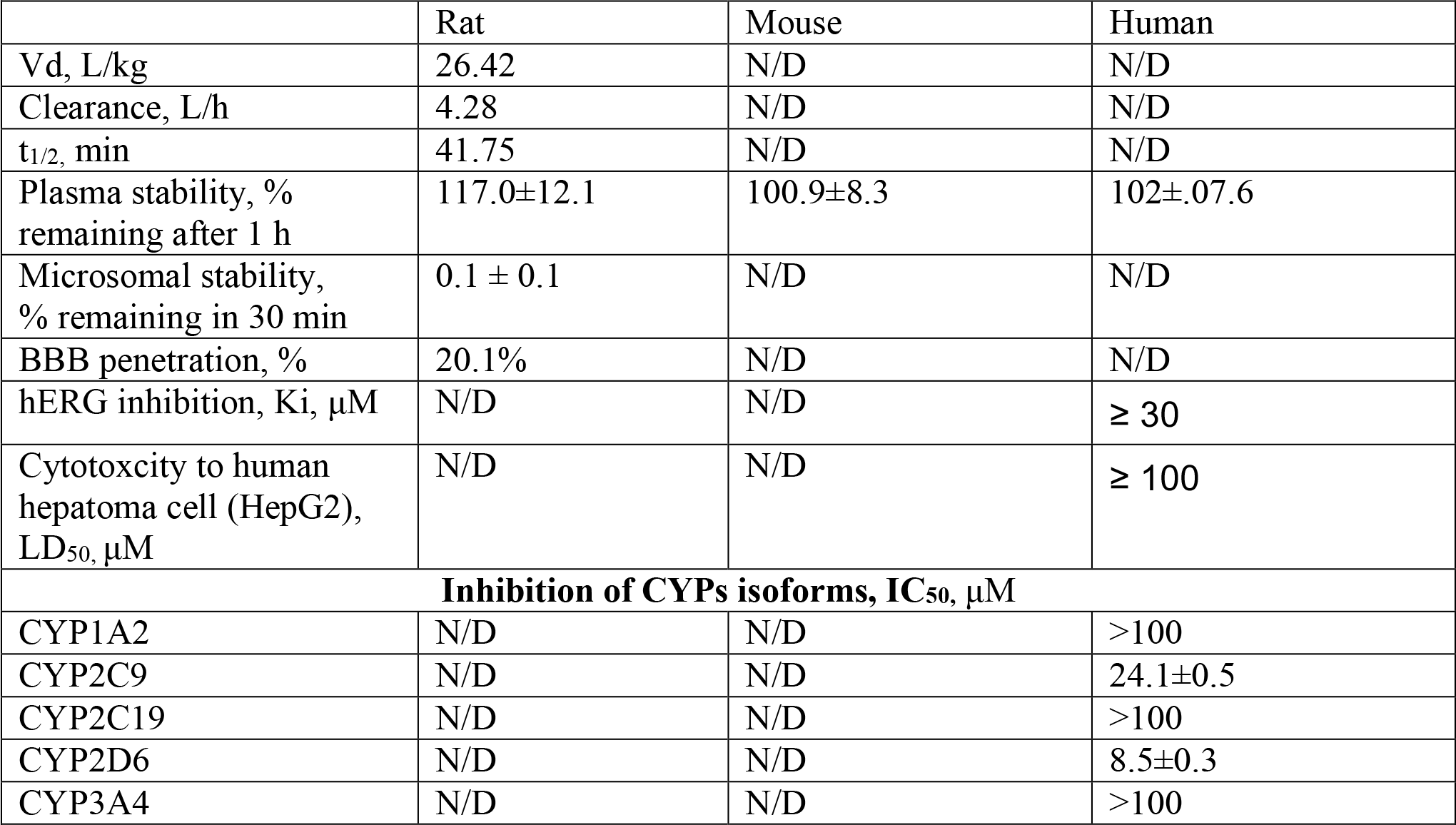
ADME properties of BT18.

|  | Rat | Mouse | Human |
| --- | --- | --- | --- |
| Vd, L/kg | 26.42 | N/D | N/D |
| Clearance, L/h | 4.28 | N/D | N/D |
| t<sub>1/2</sub>, min | 41.75 | N/D | N/D |
| Plasma stability, %<br>remaining after 1 h | 117.0±12.1 | 100.9±8.3 | 102±.07.6 |
| Microsomal stability,<br>% remaining in 30 min | 0.1 ± 0.1 | N/D | N/D |
| BBB penetration, % | 20.1% | N/D | N/D |
| hERG inhibition, Ki, µM | N/D | N/D | ≥ 30 |
| Cytotoxicity to human<br>hepatoma cell (HepG2),<br>LD<sub>50</sub>, µM | N/D | N/D | ≥ 100 |
| **Inhibition of CYPs isoforms, IC<sub>50</sub>, µM** |  |  |  |
| CYP1A2 | N/D | N/D | >100 |
| CYP2C9 | N/D | N/D | 24.1±0.5 |
| CYP2C19 | N/D | N/D | >100 |
| CYP2D6 | N/D | N/D | 8.5±0.3 |
| CYP3A4 | N/D | N/D | >100 |

BBB – blood brain barrier, hERG - **h**uman ***E**ther-á-go-go*-**R**elated **G**ene, LD50 – lethal dose 50, the dose that kills 50% of cells, Ki – constant of inhibition, IC_50_ – concentration at 50% inhibition, CYP – cytochrome P450

**Supplementary table 3.**
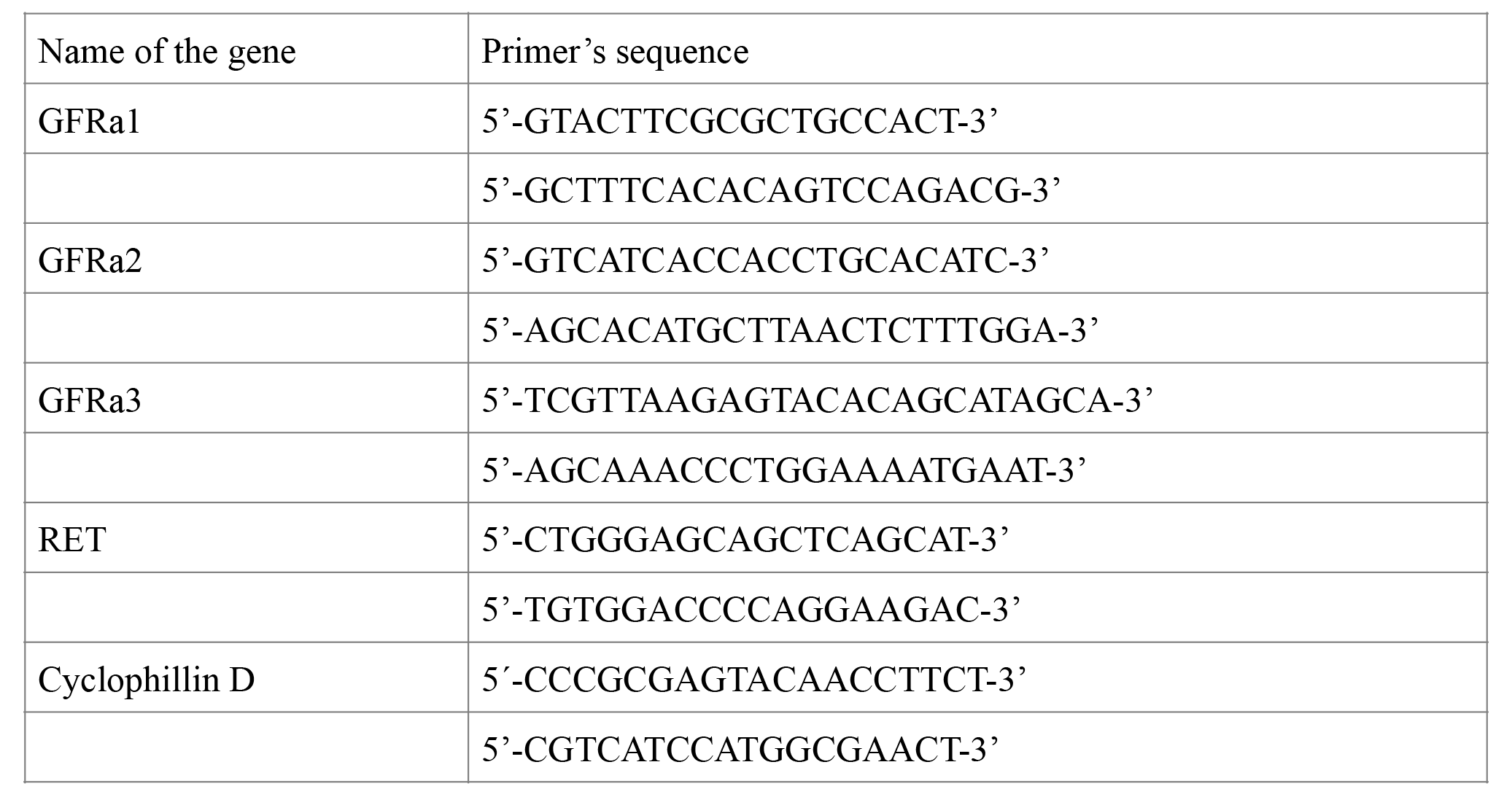
Sequence of qPCR primers for rat GFLreceptors.

### Supplemental materials and methods

#### Materials

*Cell lines*. MG87RET murine fibroblasts stably transfected with RET proto-oncogene (gift from Prof. Carlos Ibáñez). MG87TrkB murine fibroblasts stably transfected with TrkB receptor tyrosine kinase (gift from Prof. Carlos Ibáñez).

*Proteins*. GDNF, ARTN, BDNF were obtained from PeproTech Ltd. NGF was purchased from Promega. Concentration of all GFLs was checked by microBCA kit (Pierce) using BSA as a standard.

*Plasmids*. Full-length human *GFRα1* cDNA subcloned in pCDNA6 (Invitrogene; USA), full-length human *GFRα3* cDNA (was a gift of Prof. Jeff Milbrandt), full-length human *GET* (long isoform) in pCR3.1 (Invitrogen), PathDetect Elk-1 system (Stratagene, USA).

#### High throughput luciferase assays

To select GFL mimetics, we used a previously developed reporter-gene-based luciferase assay (12). The principle of this assay is described in Supplementary Fig. 1b. We used MG87 murine fibroblasts stably transfected with Pathdetect Elk-1 (Stratagene) and one of the following: RET, GFR*α*1/RET, GFR*α*1/RET. High throughput screening was performed using chemicals in a single concentration (5 μM) and single dose. The compounds were dissolved in DMSO to 5 mM concentration, transferred to 384-well storage plates (Greiner) and kept at room temperature (RT) in a desiccator before the experiment. The day before the experiment, the GFR*α*1/RET reporter cells were plated into 384-well cell culture plates (PerkinElmer) at 175 000 cell/ml density using an automatic liquid dispenser (Thermo) in 50 μM DMEM, 10% FBS, 100 μg/ml Normocin (Invivogen), 1% DMSO, 15 mM Hepes, pH 7.2. The next day, approximately 50 nl of each compound was delivered into the individual wells using a pin-tool equipped robotic workstation (Beckman Coulter). The following day, the cells were lysed using the luciferase detection reagent (SteadyGlo, Promega). Ten minutes after the reagent addition, the luminescence was measured on a TopCount luminometer (PerkinElmer).

The hit confirmation experiments, screening of focused library were performed in 96-well format in four replicas and several concentrations. For detection of luciferase activity, we used luciferase assay reagent (Promega) and did measurements on MicroBeta 2 counter (PerkinElmer).

#### Phosphorylation assays

Phosphorylation of receptor tyrosine kinase RET, TrkA, TrkB in cultured cells was assessed by antibodies against phosphorylated tyrosine residues (clone 4G10, Upstate) after immunoprecipitation of RTKs with specific antibodies targeted to RET or TrkA/TrkB, respectively. Phosphorylation of intracellular signaling proteins ERK and AKT was determined using specific antibodies against phosphorylated forms of the respective proteins. MG87RET, MG87TrkB or MG87TrkA were plated on 35 mm tissue culture dishes 1-2 days before the experiment to achieve 90-95% confluency of the cells in the day of experiments. When necessary, cells were transfected with 4 μg/well of GFR*α*1, GFR*α*3-or GFP-expressing plasmid using Lipofectamine 2000 (Invitrogen) for DNA delivery as described by manufacturer. Before the experiment, cells were starved for 4 h in serum-free DMEM containing 15 mM Hepes, pH 7.2 and 1% DMSO and stimulated with compounds or neurotrophic factors. Then cells were washed once with ice-cold PBS containing 1 mM Na_3_VO_4_ and 1 mM NaF and lysed on ice in 1 ml per well of RIPA-modified buffer (50 mM Tris-HCl, pH 7.4, 150 mM NaCl, 1 mM EDTA, 1% NP-40, 1% TX-100, 10% glycerol, EDTA-free protease inhibitor cocktail (Roche), 1 mM Na_3_VO_4_, 2.5 mg/ml of sodium deoxycholate, 1 mM NaF. Plates were incubated on horizontal shakers for 30 min with vigorous shaking. Subsequently, an 80 μl-aliquot was taken from each well and whole cell lysates prepared by mixing with equal volume of 2x Laemmli loading buffer (4% SDS, 20% glycerol, 10% 2-mercaptoethanol, 0.004% bromophenol blue, 0.125 M Tris-HCl, pH 6.8) and heating at 100°C for 10 min. Remaining cell lysates were used for immunoprecipitation of RET (or TrkA/TrkB). When immunoprecipitation was not performed, cells were plated on 48-well plates, starved and stimulated as described above, but lysed directly in 100 μl of 2x Laemmli loading buffer. To immunoprecipitate RET, TrkA and TrkB, cell lysates were incubated overnight at +4°C on the round rotator in the presence of 1μg/ml of anti-RET (C-20) or anti-Trk (C-14; recognizes both TrkA and TrkB) antibodies (Santa-Cruz Biotechnology, Inc.) and sepharose or magnetic beads conjugated with protein G. Beads were washed 3 times with 1x TBS (50 mM Tris-Cl, pH 7.4, 150 mM NaCl) with 1% Triton X-100, bound proteins were eluted by 100 μl of 2x Laemmli loading buffer, resolved on 7.5% SDS-PAGE and then transferred to a nitrocellulose membrane. The membrane was blocked for 15 min at RT by 10% skimmed milk in TBS with 0.15% of Tween 20 (TBS-T) and probed with anti-phosphotyrosine antibodies (clone 4G10, Upstate Biotechnology) diluted 1:1000 in TBS-T containing 3% skimmed milk for 2 h at RT. The membranes were washed 3 times for 5 min in TBS-T and incubated in the 1:1000 solution of secondary anti-mouse antibodies conjugated with HRP (DAKO) diluted in TBS-T containing 3% skimmed milk for 45 min at RT. Membranes were washed with TBS-T for 4x10 min. Stained bands were visualized with ECL reagent or Femto ECL reagent (Pierce) using LAS3000 imaging program. To confirm equal loading, membranes were stripped and reprobed with anti-RET (C-20) antibodies (1:500, Santa-Cruz Biotechnology, Inc.) or anti-Trk (C-14) antibodies (1:1000, Santa-Cruz Biotechnology, Inc.) in TBS-T containing 3% skimmed milk. To detect C-20 we used secondary anti-goat antibodies conjugated with HRP (1:1500, DAKO); to detect C-14 we used anti-rabbit antibodies conjugated with HRP (1:3000, GE Healthcare).

To quantitatively assess the level of RET phosphorylation we used a sandwich-ELISA based method developed by us (RET-ELISA). High-binding capacity 96-well black OptiPlates (PerkinElemer) were pre-coated overnight at +4**<u>°</u>**C with anti-RET antibodies (1 μg/ml in 1xPBS, 75μl/well) and unbounded antibodies were washed away by three changes of 1x PBS. To prevent non-specific interactions, plates were blocked in 5% BSA in 1x TBS (400 μl/well) for 3 h at RT. Afterwards, plates were washed once with RIPA buffer, incubated with cell lysates (75 μl/well) overnight at +4**<u>°</u>**C and washed three times. Then, plates were incubated with anti-phosphotyrosine antibodies (100 μl/well, clone 4G10, Upstate Biotechnology) diluted 1:1000 in binding buffer (1xTBS, 1% Triton X-100, 2% glycerol, 2% BSA) for 1-2 h at RT. Plates were washed three times with washing buffer (400 μl/well, 1x TBS, 1% Triton X-100, 2% glycerol) and incubated with secondary antibodies (100 μl/well 1:1000 anti-mouse antibodies conjugated with HRP (DAKO), diluted in binding buffer. Plates were washed three times with washing buffer. Pre-warmed Femto ECL reagent (Pierce) was added to the wells (100 μl/well). Luminescence was measured using MicroBeta 2 counter (PerkinElmer).

Whole cell lysates were resolved on 12% SDS-PAGE and transferred to nitrocellulose membranes. Membranes were blocked in 10% nonfat dry milk in TBS-T for 10-15 min and probed (with anti-phospho-Akt antibodies (Ser473, Cell Signaling) diluted 1:1000 in TBS-T containing 5% BSA overnight at +4°C or with anti-phospho-ERK antibodies (E4, Santa-Cruz Biotechnology Inc) diluted 1:500 in TBS-T containing 3% skimmed milk overnight at +4°C. Then anti-phospho-Akt-treated membrane was incubated in the 1:3000 solution of secondary anti-rabbit antibodies conjugated with HRP (GE Healthcare) and anti-phospho-ERK-treated – with anti-mouse-HRP-conjugated antibodies (1:1000, DAKO) diluted in TBS-T containing 3% skimmed milk for 45 min at RT. Membranes were washed with TBS-T 4x10 min. Stained bands were visualized with ECL reagent or Femto ECL reagent (Pierce) using LAS3000 imaging program. To confirm equal loading, membranes were stripped and re-probed with anti-GAPDH antibodies (1:4000, Millipore) diluted in TBS-T containing 3% skimmed milk. For detection we used secondary anti-mouse antibodies conjugated with HRP (1:1000, DAKO).

#### Cytotoxicity assays

To assess the ability of our compounds to influence proliferation of immortalized (MG87 murine fibroblasts) and primary non-neuronal cells (embryonic fibroblasts), we used AlamarBlue dye (Life Technologies) or CellTiterGlo (Promega). Before execution, the assay was optimized for plating cell density and length of the incubation with the dyes for each cell batch. Experiments were performed on MG87 murine fibroblasts and primary embryonic fibroblasts. To assess cell proliferation with AlamarBlue dye, cells were plated on 96-well CulturPlates (PerkinElmer) in 100 μl of DMEM, containing 10% FBS, 1% DMSO and 100 μg/ml Normocin 24 h prior assay. Compounds were dissolved in DMEM, containing 1% DMSO (final), 15 mM Hepes, pH 7.2, until 2x concentration (50 and 200 μM) and 100 μl of the solution was added to the cells. MG87 cells were incubated with compounds for another 24 h, embryonic fibroblasts – for 72 h. Afterwards, media were aspirated and replaced with phenol red-free DMEM containing 10% FBS, 100 μg/ml Normocin, and 10% AlamarBlue for 4 h. Fluorescence was detected using plate fluorimeter (Envision or Victor III Reader, PerkinElmer) with excitation wavelength 530-560 nm and emission 590 nm. Cell density was calculated from fluorescence intensity using calibrating plot.

Assessment of cell proliferation with CellTiterGlo reagent was performed according to the manufacturing instructions on 384-well plates in the same culturing conditions as described for AlamarBlue. In both cases, the signal from the wells with cells treated with vehicle (1% DMSO in DMEM, 15 mM Hepes, pH 7.2) was considered as 100%.

#### ^125^I-GDNF displacement assay

The labelling reaction was performed at RT in a final volume of 50 μl. 1 mCi Na^125^I (GE Healthcare), 1 μg GDNF protein, 0.5 U lactoperoxidase (Sigma, St. Louis, MO), 2 μl 0.03% H_2_O_2_ were mixed in 0.1 M sodium phosphate buffer, pH 6.0, and incubated for 10 min at RT. Subsequently, an additional 2 μl of H_2_O_2_ was added and incubation continued for another 10 min. The reaction was terminated by adding 3 volumes of buffer containing 0.42 M NaCl, 0.1 M NaI, and 0.1 M sodium phosphate buffer, pH 7.5, and 22 μl of 2.5% BSA (Sigma), in 0.1 M sodium phosphate buffer, pH 7.5. Free iodine was separated from GDNF using PD-10 desalting columns (GE Healthcare). To assess competition of BT18 and ^125^I-GDNF, HEK cells were transfected with RET-and GFR*α*1-expressing plasmids at a 1:1 ratio or GFR*α*1-expressing plasmid 2 days before assay. On the day of assay, cells were incubated on ice in blocking solution (DMEM, 15 mM Hepes, pH 7.4, 1% DMSO, 5% Sea Block (Pierce)) for 30 min. Afterwards, cells were treated with 5-100 μM BT18 or 5 nM unlabelled GDNF and 50 pM of ^125^I-GDNF dissolved in blocking solution and incubated for 2 h on ice, washed 4 times with binding buffer without Sea Block and lysed with 1 M NaOH. Lysates were transferred to vials and counted on a Wallac Gamma Counter(Wallac/LKB). The assay was performed in quadruplicates. Results are presented as Mean±SEM of five independent experiments.

#### Neurite outgrowth from DRG neurons

DRG ganglia were dissected from P1-P3 newborn mice in PBS under a dissection microscope. Ganglia were cleaned from nerve fibers, transferred to a centrifuge tube and washed twice with HBSS. To prepare dissociated neuronal cultures, ganglia were incubated in trypsin (10 mg/ml) for 20 min at 37°C, followed by DNase I treatment for 5 min. DRGs were washed twice with trituration solution (HBSS, DNase I) and triturated with siliconized glass Pasteur pipette in a fresh portion of warm HBSS. Cells were precipitated by centrifugation at 1000 rpm in Eppendorf centrifuge (5702), washed 2 times with growth media (Neurobasal media, B27 serum supplement, 1 mM L-glutamine, 100 μg/ml primocine) and plated on glass cover slips or cell culture dishes coated at first with poly-ornithine (Sigma-Aldrich) (overnight at +4°C) and subsequently with 1 mg/ml of laminin (Sigma-Aldrich) (at least 4 h at +37°C) into the media containing different concentrations of BT18 or 30 ng/ml NGF or 30 ng/ml ARTN (positive controls). To assess neurite outgrowth at 15-20 h post plating, cultures were fixed in 4% PFA in PBS for 15 min at RT, washed three times with PBS, probed with 1:500 rabbit anti-PGP9.5 antibodies (Enzo) and visualized with anti-rabbit secondary antibodies conjugated to Alexa488 (Invitrogen). The number of neurons with neurites at least twice as long as the cell body were counted under a fluorescent microscope (Olympus IX70) and normalized to the total number of neurons. Each compound in each concentration was tested in quadruplicate. Experiments were repeated twice on independent cultures.

#### ADME-Tox profiling

ADME profiling was performed by CRO Cyathus Exquirere (Milan, Italy). To assess BT18 pharmacokinetics, 10 mg/kg of BT18 were intravenously injected into nine male Sprague Dawley rats. Plasma and brain samples were collected 1, 3 and 6 h later. Plasma stability of BT18 was assessed after 1 h incubation with human, rat and mouse plasma, using verapamil as internal standard and lidocaine and M7319 (Sigma) as reference compounds. Metabolic stability of BT18 was measured after 30 min incubation with 0.5 mg/ml of rat liver microsomes (Xenotech) using verapamil as internal standard and 7-ethoxycoumarin and propranolol as reference standards. In all above-mentioned cases, samples were analyzed by chromatography on an UPLC system integrated with a Premiere XE Triple. To assess cardiac safety of BT18 we measured its K_i_ in competition assay with radiolabelled ligand of the hERG (human Ether-**à**-go-go-related gene) [^3^H]-astemizole in crude cell membrane fraction of HEK293 cells stably transfected with the corresponding gene. BT18 toxicity towards HepG2 cells was determined using the MTT assay. Inhibition of the most important drug-metabolizing isoforms of CYP (CYP1A2, CYP2C9, CYP2C19, CYP2D6 and CYP3A4) was measured using a fluorescent assay with specific substrate for each isoenzyme. The IC_50_ was determined using a four-parameter curve fit. Furafylline, sulfaphenazole, tranylcypromine, quinidine and ketoconazole were used as reference compounds for CYPs listed above.

#### Immunohistochemistry

Sections of sciatic nerves and DRGs from rats with spinal nerve ligation and treated with BT18 or vehicle were probed, after deparaffinization and acidic or basic antigene-retrieval using standard IHC protocols, with antibodies against Na_V_1.8 (1:1000, Abcam, UK), IB-4 (1:200, Griffonia), P2X3 (1:1000, Neuromics), CGRP (1:10000, Peninsula Laboratory), NPY (1:10000, Peninsula Laboratory), pERK1/2 (1:300, Cell Signaling), pS6 (1:300, Cell Signaling), pan-neuronal marker PGP9.5 (1:500 Abcam) and nuclear dye. The secondary antibodies were conjugated with fluorophors (Alexa 488, Alexa 568 and Alexa 647, Invitrogen) or horseradish peroxidase (Sigma) in combination with ABC and DAB staining kits (VectoLabs) according to the manufacturer’s instructions. Whenever possible, double labelling with antibodies against the specific markers and PGP9.5 was performed. Nuclei were visualized using hematoxilin or DAPI dyes. To prevent the bleaching of the dyes, microscopic slides were mounted using Immu-Mount or Percoll, images were taken with fluorescent or bright-field microscopes (Carl Zeiss, Axio Imager.M2) and analyzed using MatLab R2012a software. In DRGs, the number of the cells positive for specific marker was normalized to the total number of the neurons (identified by PGP9.5 staining or by morphology). In sciatic nerve the intensity of Na_V_1.8 staining only in PGP9.5-positive regions was assessed. Statistical analysis was performed using data from 3-5 individual animals.

#### qPCR

Expression of GFL receptors (GFR*α*1-3, RET) in DRGs of animals with SNL receiving vehicle, BT18 or ARTN (0.5 mg/kg) were analyzed by qPCR. RNA isolation and DNA digestion of the samples was performed using NucleoSpin^®^ RNA kit (Macherey-Nagel Inc.) according to the manufacturer’s instructions. RNA concentration was measured with NanoDrop (ND 1000); cDNA was synthesized by Maxima H minus reverse transcriptase (ThermoScientific Inc.) in total volume 20 μl as described by manufacturer. 15 μl of the mixture containing 100 ng of RNA, 100 pmol of oligo(dT)^15-18^ (Promega, USA; Oligomer, Finland), 0.5 mM dNTP (ThermoScientific) were incubated at 65°C for 5 min to remove possible secondary structure and melt GC-rich regions, and chilled on ice and combined. Afterwards, reverse transcriptase buffer (50 mM Tris-HCl, pH 8.3, 75 mM KCl, 3 mM MgCl_2_ 10 mM DTT, final concentrations), 20 U of RiboLock RNase Inhibitor and 100 U of Maxima H Minus Reverse Transcriptase (all from ThermoScientific) were added. The mixture was incubated for 30 min at 50**<u>°</u>**C and reaction was stopped by heating at 85**<u>°</u>**C for 5 min. qPCR was performed using LightCycler^©^ SYBR Green I Master Mix (Roche Diagnostics) according to the manufacturer’s instructions in a total volume of 10 μl. The reaction mixture contained 0.25 μ1 of cDNA, 1xSybrGreen I Master Mix, 2.5-10 pmol of forward and reverse primers. Sequences of the primers are presented in Supplementary Table 3. Relative expression of the target genes was determined by the 2^-Δ ΔCT^ method; as a normalization standard cyclophilin A was used. Statistical analysis was performed using data from 4-5 individual animals per group.

## Abbreviations

ADME: absorption, metabolism, distribution, excretion
ADME-Tox: ADME-toxicity in pharmacokinetics
Akt: protein kinase B (PKB)
BDNF: Brain-derived neurotrophic factor
BL: baseline
ARTN: artemin
CTR: control
Elk1: transcription factor activated by MAPK
ERK: extracellular signal-regulated kinase
GAPDH: glyceraldehyde-3-phosphate dehydrogenase
GDNF: glial cell line-derived neurotrophic factor
GFL: glial cell line-derived neurotrophic factor family ligand
GFR*α*: GDNF family receptor-*α*
GPI: glycosylphosphatidylinositol (anchors GFR *α* receptors to the membrane)
IP: immunoprecipitation
MAPK: mitogen-activated protein kinase
NGF: nerve growth factor
NP: neuropathic pain
NRTN: neurturin
p-: phosphorylated form of…
PLC *γ*: phospholipase C gamma
PI3K: phosphoinositide 3-kinase
PSPN: persephin
pY: phosphotyrosine
RET: RTK rearranged during transfection
RTK: receptor tyrosine kinase
Src: (proto-oncogene) tyrosine-protein kinase
WB: Western blotting

